# Physical activity improves sarcopenia in a murine model by enhancing the proliferative potential of muscle stem cells, oxidative capacity of mitochondrial enzymes and expression of Sestrins

**DOI:** 10.1101/811638

**Authors:** Masroor Anwar, Saumya Ranjan Mallick, Daizy Paliwal, Shashank Sekhar, S.K. Panda, Sharmistha Dey, A.B. Dey

**Affiliations:** Department of Geriatric Medicine, All India Institute of Medical Sciences, New Delhi, India; Department of Pathology, All India Institute of Medical Sciences, New Delhi, India; Department of Biophysics All India Institute of Medical Sciences, New Delhi, India

**Author notes:** **Address for Correspondence**. Dr. Aparajit B Dey, Department of Geriatric Medicine, All India Institute of Medical Sciences, Ansari Nagar, New Delhi - 110029, India; Dr. Sharmistha Dey, Department of Biophysics, All India Institute of Medical Sciences, Ansari Nagar, New Delhi - 110029, India.

**Keywords:** Ageing, Sarcopenia, Muscle stem cells, Physical activity, Sestrins

## Abstract

Sarcopenia is a major health issue in old age. Underlying molecular mechanisms in its genesis remain unclear and optimal animal models are yet to be established. A novel animal model was developed to identify structural and functional changes in skeletal muscles from sarcopenia. The influence of physical activity on animals of the sarcopenic model with respect to the expression of anti-oxidant proteins, mitochondrial oxidative capacity, and effect on muscle stem cells (MuSCs), were assessed. Male C57Bl/6 mice of different age groups were recruited: Y-Cntrl (young-control), A-Cntrl (aged-control), SAR-model and SAR-INT. SAR-model group was kept in a retrofitted confined cage and was fed with customized protein-restricted diet (14% protein), which is considered as a sarcopenic featured model. SAR-INT served as the intervention group. Three parameters, namely, muscle mass, grip strength, and physical endurance, were used to confirm the sarcopenic state. All physical parameters deteriorated most in SAR-model and it improved in the SAR-INT group. The impact of physical activity on the SAR-INT group was also evidenced by the improved proliferative potential of MuSCs determined by flow cytometric analysis. Compared with the SAR-model, the SAR-INT exhibited significant improvement in the oxidative capacity of mitochondrial enzymes and increased expression of anti-oxidant proteins, sestrins. In conclusion, physical activity improved physical parameters, MuSC proliferative potential, mitochondrial enzyme oxidative capacity and sestrin expression in sarcopenic animals. Hence, SAR-model in mice can serve as a novel sarcopenic model, physical activity provides scope for improvement in sarcopenic population and sestrin molecule can have a potential implication on sarcopenia.

## 1. Introduction

Skeletal muscle is one of the most dynamic and plastic tissues, accounting for almost 40% of total body weight and 50-75% of all the proteins in the body(Wolfe, 2006). Skeletal muscles contribute to over 25%ofprotein synthesis in the body (Liu, Mac Gabhann, & Popel, 2012). Moreover, skeletal musclesare essential for locomotion and have important roles inthe metabolism of several nutrients. Thus, changes in skeletal muscle mass and composition, such as the changes that occur in sarcopenia-cachexiaandatrophy-adversely affect health, well-being and the ability to carry out daily life activities.

Ageing has a significant effect on the development and function of skeletal muscles. After maximum muscle size and strength are attained in young adulthood, muscle area starts to decreasein the fourth decade of life(Short, Vittone, Bigelow, Proctor, & Nair, 2004). The strength and quality of leg muscles normalized to fat-free massdecreases with age(Nair, 2005).Muscle massdecreases at an annual rate of 1–2% after the fifth decade, andstrength declines by 1.5% between ages 50 and 60 and thereafter by 3% annually(von Haehling, Morley, & Anker, 2010). Limitedmobility in advancing years is an outcome of the aforementioned biological processes, frequently termed sarcopenia(Rosenberg, 1989). Sarcopenia has been defined as an age-related involuntary loss of skeletal muscle mass and strength(Walston, 2012). Since October 2016, sarcopenia has been included in theInternational Classification of Diseases, Tenth Revision, Clinical Modification (ICD-10-CM), as a diagnosis in clinical practice(Falcon & Harris-Love, 2017). Sarcopenia has been associated with increased vulnerability, decreased physical function andadverse outcomeswhen combined with acute illness and stress(Walston, 2012).Sarcopenia can result fromthe disuseof musclesdue to being bedbound, endocrine disorders, chronic diseases, inflammation, insulin resistance, and multiple nutritional deficiencies.

The consensus diagnostic criteriafor sarcopenia haveonly recently been establishedfor European and Asian populations. Thus, the exact estimates ofthe prevalenceofsarcopeniain humansarenot well established, though several estimates and projections exist intheliterature(Janssen, 2011). Sarcopenia is also a common feature in the syndrome of frailty, another clinical state in geriatric practice.The management of sarcopenia remains challenging due to a lack of clear understanding of the origin and biological processes that lead to the condition. Research on muscle stem cells (MuSCs) has provided some hope of reversing the unintentional decrease in muscle mass that occurs in sarcopenia. Adult MuSCs, originally called satellite cells, are essential for muscle repair and regeneration throughout life (Bengal, Perdiguero, Serrano, & Muñoz-Cánoves, 2017). The regeneration of skeletal muscle depends on a population of adult stem cells (satellite cells) that remain quiescent throughout life. The regenerative functions of satellite cellsdecline with age(Sousa-Victor et al., 2014). Failure of the regeneration machinery in sarcopenic muscle to replace damaged myo-fibrescan be one of the major causes of physical incapacitation and loss of independence in olderindividuals(Ibebunjo et al., 2013).

It is commonly observed that physical exercise improves muscle mass and strength in young individuals. In older persons, physical activity improvesfitness, mobility and independence, which isthe basis of physical therapy as a treatment modality in diverse disease conditions.Despite common knowledge of thebenefits of physical activity, themajority of olderpeople live a sedentary life, which may be one of the reasons for frailty and sarcopenia in old age.

Oxidative stress induced by reactive oxygen species (ROS) may have a role in genesis of sarcopenia. ROS play a critical role in loss of skeletal muscle (Xie et al., 2015) in old age. Oxidative stressis one of the causes of apoptosis of progenitor and mature skeletal muscle cells (Bellanti et al., 2018). It has also been reported that exercise upregulates anti-oxidant defences in skeletal muscle (Ji, 2007) and that anti-oxidant genes are upregulated after moderate exercise (Gomez-Cabrera, Domenech, & Viña, 2008). Considering the role of oxidative stress conditions in the genesis of sarcopenia, several antioxidant molecules were explored. Sestrin proteins appeared to be promising in animal models of sarcopenia. The Sestrin family of proteins (Sestrin 1 and Sestrin 2) has been studied for their roles in skeletal muscle biology. Sestrin, as an antioxidant molecule, has shown altered expression in frail patients and plays a protective role in stressful conditions (Rai et al., 2018). Exercise is a powerful non-pharmacological tool that can induce the renewal of the satellite cell pool in skeletal muscles (Kadi & Ponsot, 2010).

Skeletal muscle is not a stable tissue, as satellite cells are constantly recruited during normal daily activities. MuSCs have shown rejuvenation properties in cases of stress and injury. We developed a sarcopenic animal model similar to sarcopenia of ageing by restricting dietary protein and physical activity. In this model, structural and functional changes in skeletal muscles and characteristics of MuSCs in aged animals were studied. Parameters for assessment of sarcopenia were similar to those used in clinical practice: body weight, grip strength and gait speed. Molecular aspect of functional improvement and enhancing MuSC activity were studied through physical activity intervention in the developed murine model. The role of antioxidant protein, Sestrin, in the modulation of MuSC regenerative potential was also evaluated. This is the first study which reported the role of sestrin protein in the newly developed sarcopenic mouse model

## 2. Results

Eighty male C57BL/6mice of six to eight weeks age were included in this prospective study. These animals were assigned to the following groups: young control (Y-Cntrl), sacrificed at 14 months; aged control (A-Cntrl), sacrificed at 21 months; protein-deficient model (SAR-model) sacrificed at 21 months; and post-intervention SAR-model (SAR-INT) sacrificed at 24 months. The goal of the experiment was to develop an animal model with features of sarcopenia through protein deficiency and sedentary life style, which mimic older sarcopenic human beings and will serve as a platform for intervention such as physical exercise.

### 2.1 Physiological parameter tests

#### 2.1.1 Body weight

To analyse differences in animal body weight, the mean body weight of the animals in all four groups was calculated. A significant decline in mean body weight was observed in all animal groups. An 8.6% decrease in mean body weight was observed in SAR-model animals compared to A-Cntrl animals. Analysis of the effect of physical activity on body weight revealed no significant difference between the SAR-model and SAR-INT groups. The mean body weight of the A-Cntrl group was higher than that of the Y-Cntrl group, indicating probable fat accumulation with ageing (Figure 1a).

**Figure 1.**
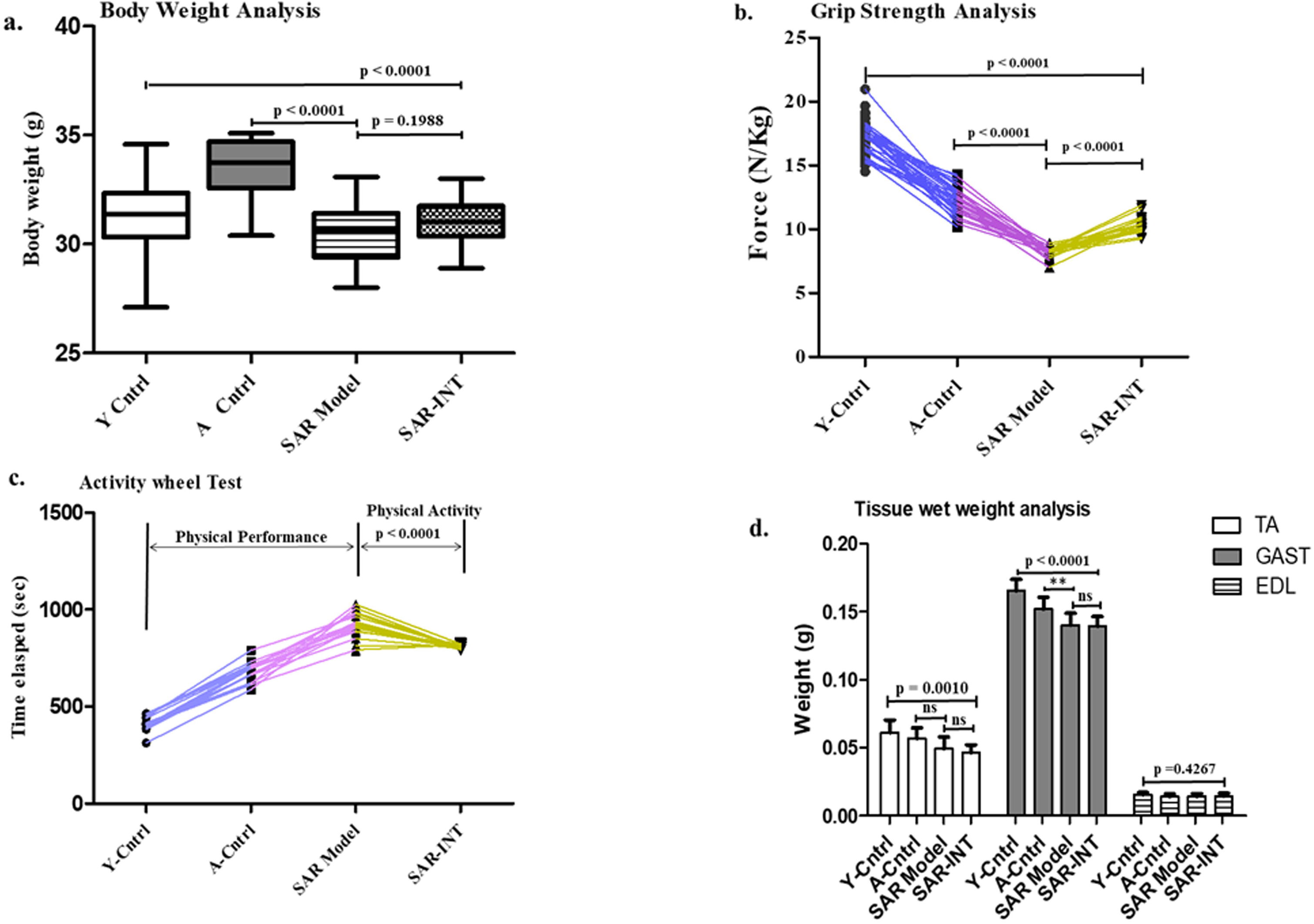
Phenotypic assessment: (a) Animal Body Weight, (b) Grip Strength Test, (c) Activity Wheel Running Test, (n=20-30) and (d) Tissue Wet Weight analysis (n=6-11) of all the experimental groups: Y-Cntrl, A-Cntrl, SAR-model and SAR-INTm(p<0.0001).

#### 2.1.2 Grip strength

Grip strength was expressed as a measure of force in N/kg and reflected the changes. The highest values were recorded in the young animals. Grip strength gradually declined with increasing age. Compared to the A-Cntrl group, the SAR-model group exhibited a nearly 32% decrease in grip strength, showing the effect of low protein dietary intake and sedentary lifestyle. Compared with the SAR-model group, the SAR-INT group showed 25% increase in grip strength, which can be considered as functional improvement after physical activity (Figure 1b).

#### 2.1.3 Time elapse analysis by activity wheel test for physical performance

Time elapse analysis was performed using an established protocol. The test measured the ability of mice torun on an activity wheel to assess the physical performance of the animal. Analysis was performed for only three groups (Y-Cntrl, A-Cntrl, and SAR-Model) to evaluate the baseline for each experimental group. The time required to complete a 100-metre run was found to proportionally increase with increasing age (Y-Cntrl vs. A-Cntrl), with a longer time indicating a lower gait speed and endurance capacity. The results of the time elapse analysis showed 28% increase in the time elapsed in the SAR-model group compared to the A-Cntrl group. After physical activity, the SAR-INT group showed a significant 14% improvement compared to the SAR-model group (Figure 1c).

### 2.2 Histology, morphometry and enzyme histochemistry

#### 2.2.1 Wet Tissue weight assessment

The wet weight of tissues was measured to analyse the changes that occurred at the tissue level. Tibialis anterior (TA), gastrocnemius (GAST) and extensor digitorum longus (EDL) muscles were weighed for this experiment. Compared with the A-Cntrl group, the SAR-model group showed significant (7.89%) decrease in the wet weight of GAST tissue and13.5% decrease in TA weight. However, no significant improvement in the wet weight of the examined muscles was observed in the SAR-INT group compared with the SAR-model group. The weight of GAST tissue was similar in the in the SAR-INT group and the SAR-model group, with no noticeable difference. No significant change was observed in EDL muscle in any of the groups (Figure 1d).

#### 2.2.2 Mitochondrial enzyme expressions with physical activity in murine model featuring sarcopenia by histological staining

Adenosine triphosphatase (ATPase) staining was carried out to analyse the changes in the cross-sectional area (CSA) of muscle fibres. The area expressed in μm^2^ was calculated for type II fibres. A 10% reduction in CSA was noted in the SAR-model group compared to the A-Cntrl group. An 11.3% increase in the CSA was observed in the SAR-model group compared to the SAR-INT group (Figure S1). Semi-quantitative analysis of the intensity of ATPase enzyme expression in type II fibres was performed for all four experimental groups. The lowest ATPase enzyme expression was noted in the SAR-model group, with a decrease of 38% compared to that of the A-Cntrl group. An 18.9% increase in enzyme expression was observed in the SAR-INT group compared to the SAR-model group. No significant difference in enzyme expression was observed between Y-Cntrl and A-Cntrl animals (Figure 2a).

**Figure 2.**
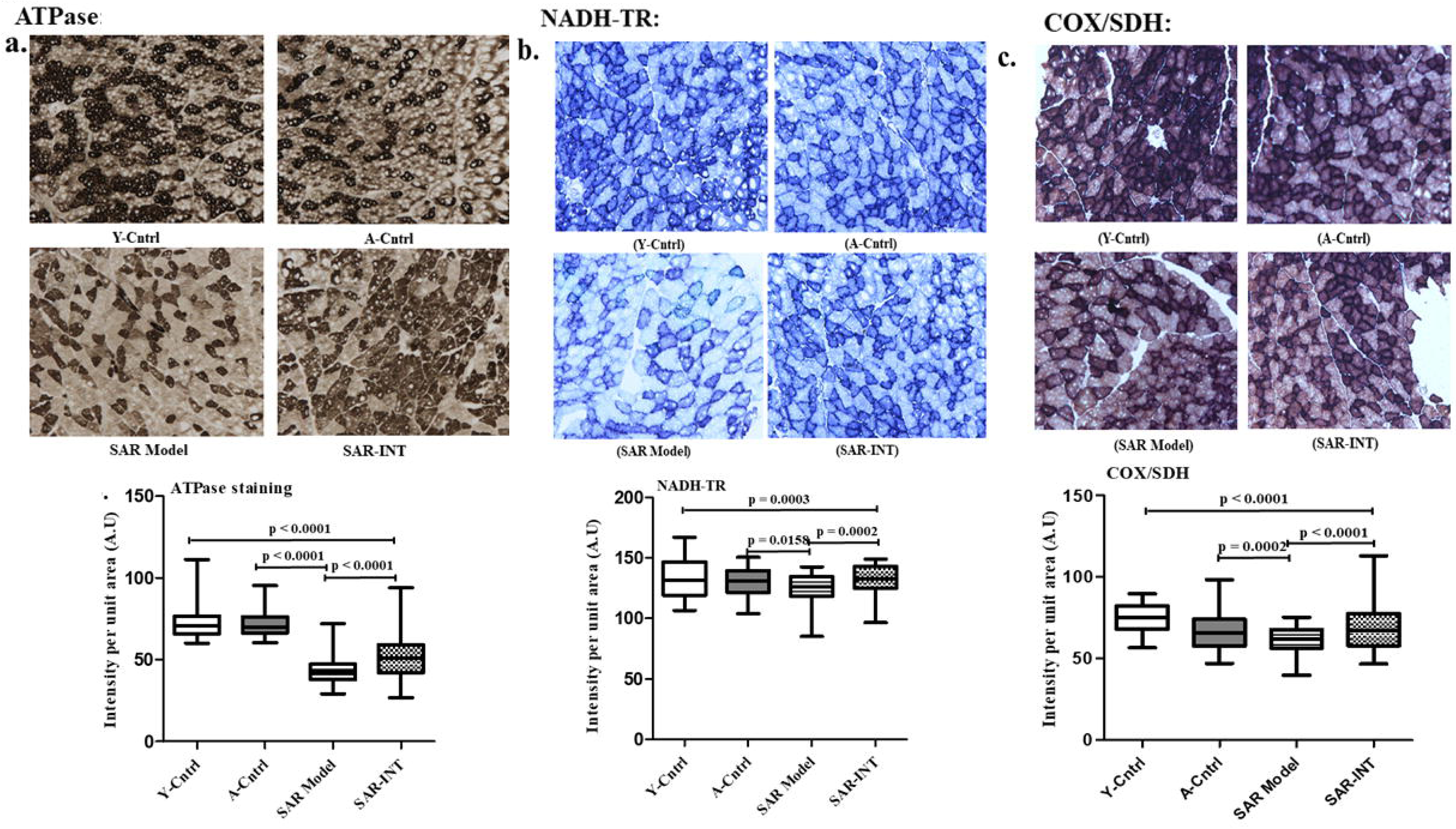
Enzyme staining images & Differential expression of mitochondrial enzyme intensity in Whisker Plot (n=3-5): (a) ATPase, (b) NADH-TR, (c) COX/SDH.:

The oxidative capacity of the mitochondrial enzymes in the skeletal muscles of experimental animals was evaluated by the relative quantification of staining for the enzyme nicotinamide adenine dinucleotide tetrazolium reductase (NADH-TR). The intensity of the SAR-model group was lower than that of the age-matched control (A-Cntrl). Compared with the A-Cntrl group, the SAR-model group exhibited 3.4% decrease in NADH-TR staining. In contrast, 5.9% increase in enzymatic intensity was observed in the SAR-INT group compared to the SAR-model group (Figure 2b).

Mitochondrial respiratory enzyme function was assessed with COX/SDH staining. Variation in the shade of the brown precipitate among the different experimental groups is shown in Figure 2c. The highest respiratory enzyme function was observed in the Y-Cntrl group. Compared to the age-matched control (A-Cntrl), the SAR-model group demonstrated 8.9% decrease in enzyme function. The SAR-INT group showed 13.1% higher staining intensity than the SAR-model group.

### 2.3 Double Immunofluorescence Assay

To locate and quantify MuSCs, a double Immunofluorescenceassay (IFA) was performed using PAX 7 (a canonical marker expressed in MuSCs) and laminin (a basement membrane protein that forms the outer membrane of muscle fibres). Haematoxylin and eosin staining was performed to visualize the tissue morphology (Figure S2). Images showed tissue with intact membranes and nuclei at the periphery of the membrane. The IFA analysis of the MuSCs from young and aged animals along with the negative control is presented in Figure 3a in panel 1, 2 and 3 respectively. MuSCs were found to be located between the basement membrane and sarcolemma in skeletal muscle tissue. The quantification of Pax 7-positive cells and laminin-positive cells with DAPI-labelled nuclei showed that the number of double-positive cells per 100 fibres decreased by 20% in the A-Cntrl group compared to the Y-Cntrl group (Figure 3b). In addition, when the number of nuclei per 100 fibres in the Y-Cntrl and A-Cntrl groups was calculated and compared, the observed decrease was not significant (Figure 3c).

**Figure 3.**
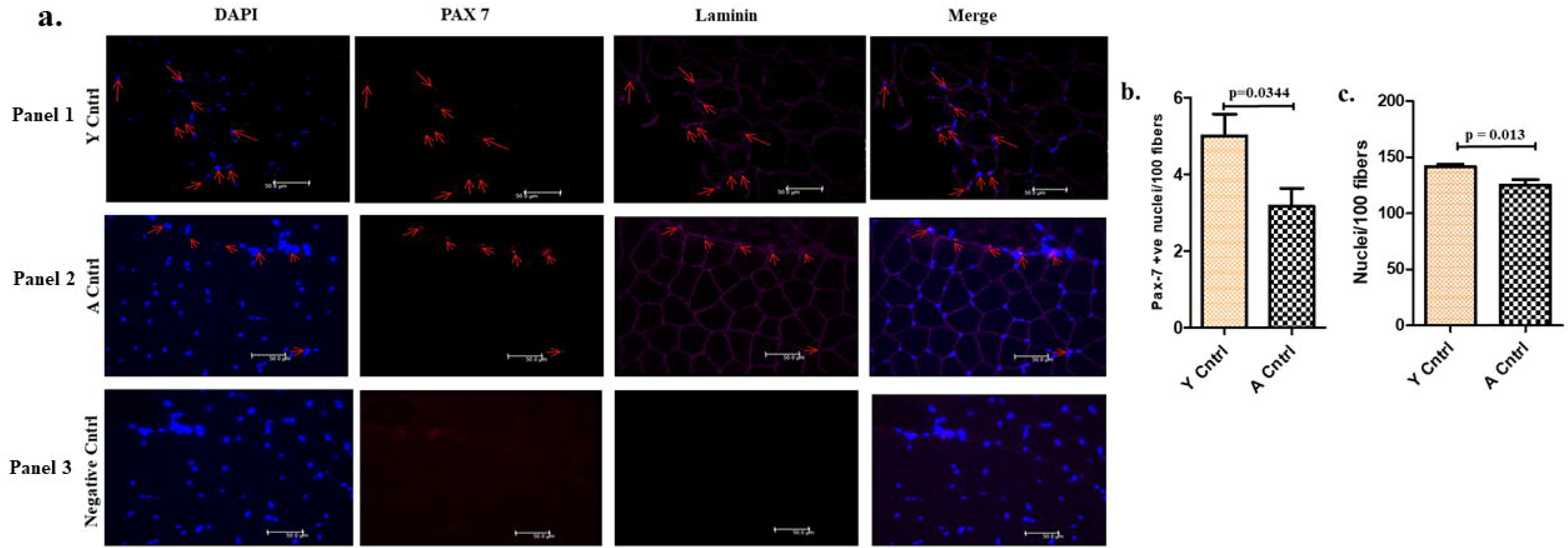
Morphometric characterization of skeletal muscle tissue of young and aged for muscle stem cells quantification: (a) Double-immunofluorescence images stained for Pax 7, Laminin along with DAPI to stain nuclei in panel 1, Y-Cntrl, panel 2, A-Cntrl and panel 3 negative-Cntrl. (b) Comparative decline in young and aged in Pax 7 positive nuclei/100 fibers (c) nuclei/100 fiber count (n=3).

### 2.4 Flow Cytometry Analysis: Proliferative potential of MuSCs with physical activity in sarcopenic model

MuSCs were isolated using the magnetic-activated cell sorting (MACS) technique. Figure 4a and 4b illustrating unstained cells and Y-Cntrl cells. A 50% decrease in the number of MuSCs positive for both (quadrant I) alpha Int-7 and Ki-67was observed in the SAR-model group compared to the A-Cntrl group. This result reflected the decreased proliferation potential of the MuSC population in the SAR-model group compared to the A-Cntrl group (Figure 4c and d). Physical activity had a beneficial effect on the MuSC population in the SAR-INT group. The number of double-positive MuSCs in the SAR-INT and SAR-model groups was compared. A three fold increase in the MuSCs with proliferative potential was observed in the SAR-INT group compared to the A-Cntrl group (Figure 4e). Notably, this increase in double-positive cells indicates that the rejuvenation potential of MuSCs in the SAR-model with sarcopenia was rescued.

**Figure 4.**
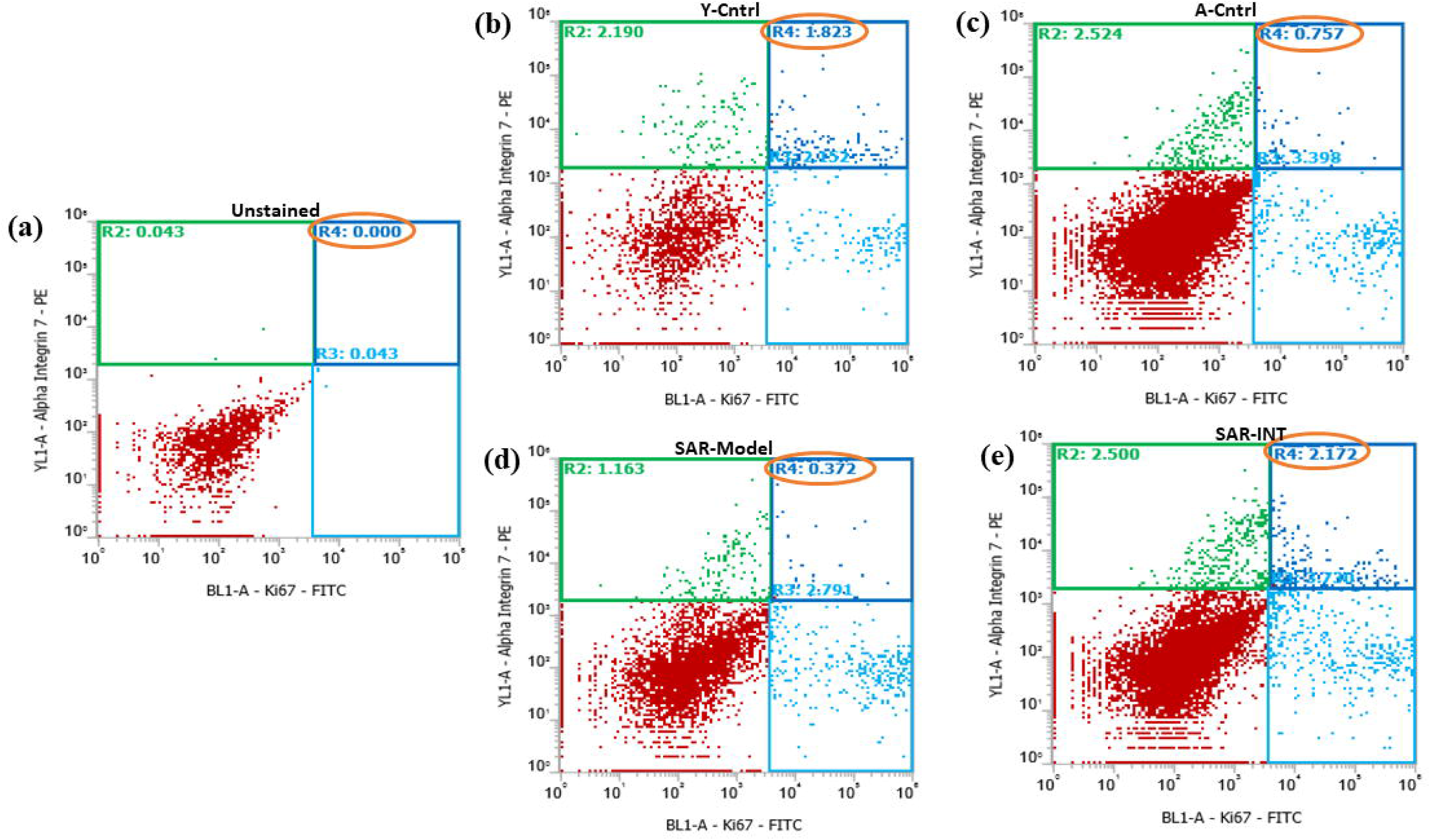
Dot plot for Flow Cytometry: X-axis representing PE-Conjugated α-Integrin 7, Y-Axis having FITC-Conjugated Ki-67(n=3). (a) Unstained cells (b) Y-Cntrl group (c) A-Cntrl (d) SAR-Model and (e) SAR-INT

### 2.5 Expression of Sestrin proteinswith physical activity in sarcopenic model group

The concentration of the antioxidant protein, Sestrin, was measured using Surface Plasmon Resonance (SPR) technology. Differences in the relative expression profile of Sestrins were observed by running samples on an immobilized chip. The response unit (RU) values of the recorded immobilized antibodies were 3940 RU and 4850 RU, where 1 RU corresponds to 1 pg/mm^2^. The Sestrin 1 concentration was 2.9% higher in the SAR-INT group than in the SAR-model group (Figure 5a), while it was 1.7% lower in the SAR-model group than in the A-Cntrl group. Sestrin 1 showed a similar expression profile across the groups. The Sestrin 2 concentration was significantly higher in the SAR-INT group than in the A-Cntrl and SAR-model groups (Figure 5b). The expression level of Sestrin 2 in the SAR-INT group was 1.65 ng/μl (28%) higher than that in the SAR-model group. The expression level of Sestrin 2 in the SAR-model group was 14% lower than that in the A-Cntrl group.

**Figure 5.**
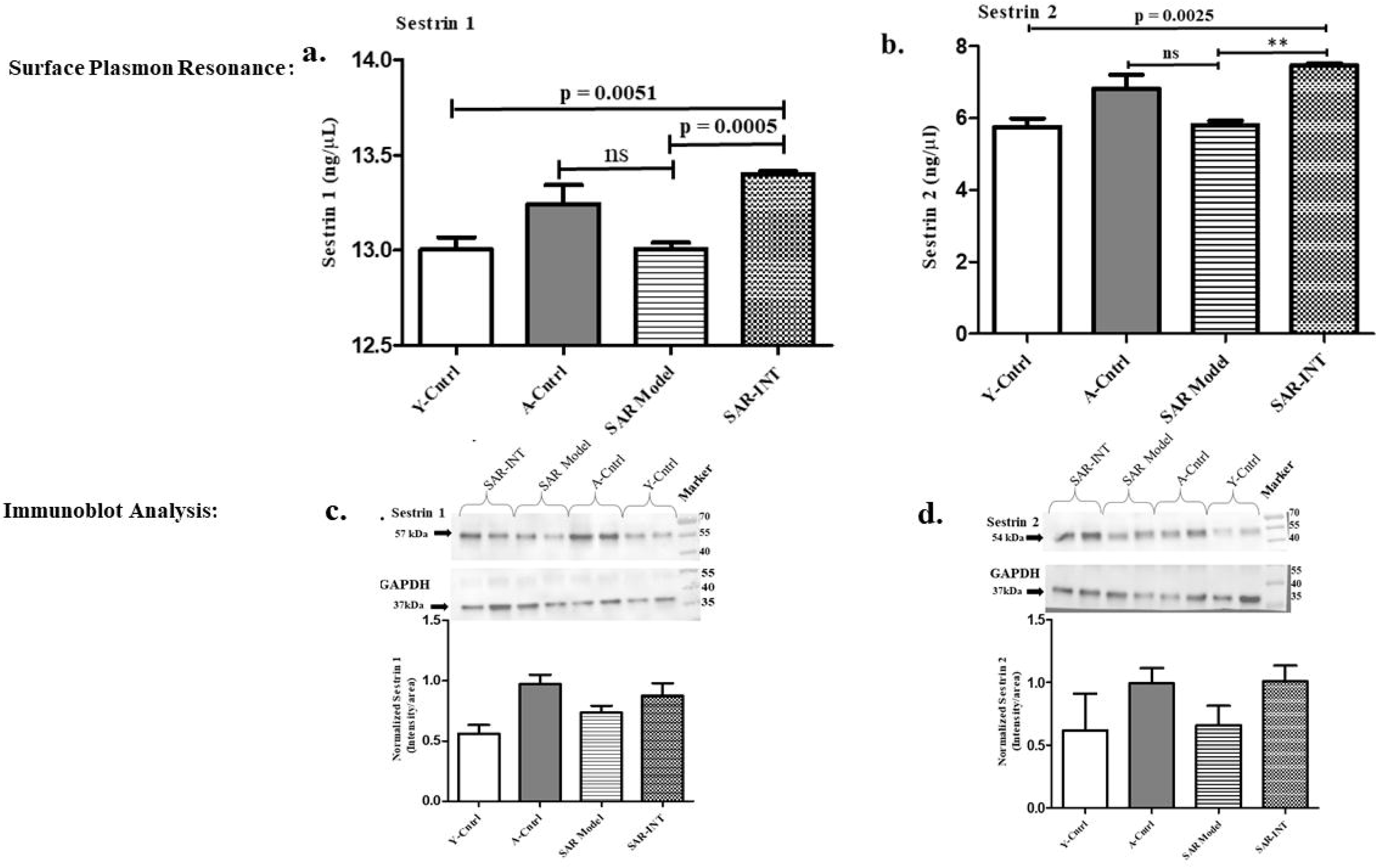
Quantification of Sestrin 1 and Sestrin 2: (a & b) Surface Plasmon Resonance and (c & d) western blot lower panel (n=3-4).

### 2.6 Western blot Analysis for validation of Sestrin expression across the groups

The quantified SPR data were validated by Western blot analysis. Cell lysates from all four animal groups were analysed for differential expression of Sestrin 1 and Sestrin 2 (Figure 5c and d). The band intensities of normalized Sestrin showed increased expression in the SAR-INT group compared with the SAR-model group for both sub-units 1 and 2. The band intensity of the SAR-model group was found to be weaker than that of the A-Cntrl group, representing a decline in concentration with age. In the SAR-INT group, the band intensity of Sestrin 2 was found to be significantly elevated compared with that of Sestrin 1, which is clearly consistent with the SPR findings. Here, glyceraldehyde-3-phosphate dehydrogenase (GAPDH) was used to normalize the band intensities of both protein subunits.

### 2.7 Differential expression of superoxide dismutase (SOD) enzyme in the experimental groups

Cellular SOD quantified through the spectrophotometric analysis showed a decline of 22% in A-Cntrl when compared to Y-Cntrl. Though there was no significant difference in SOD level between SAR-model and SAR-INT, but there was increase in the expression of SOD in SAR-INT compare to SAR-model (Figure 6).

**Figure 6.**
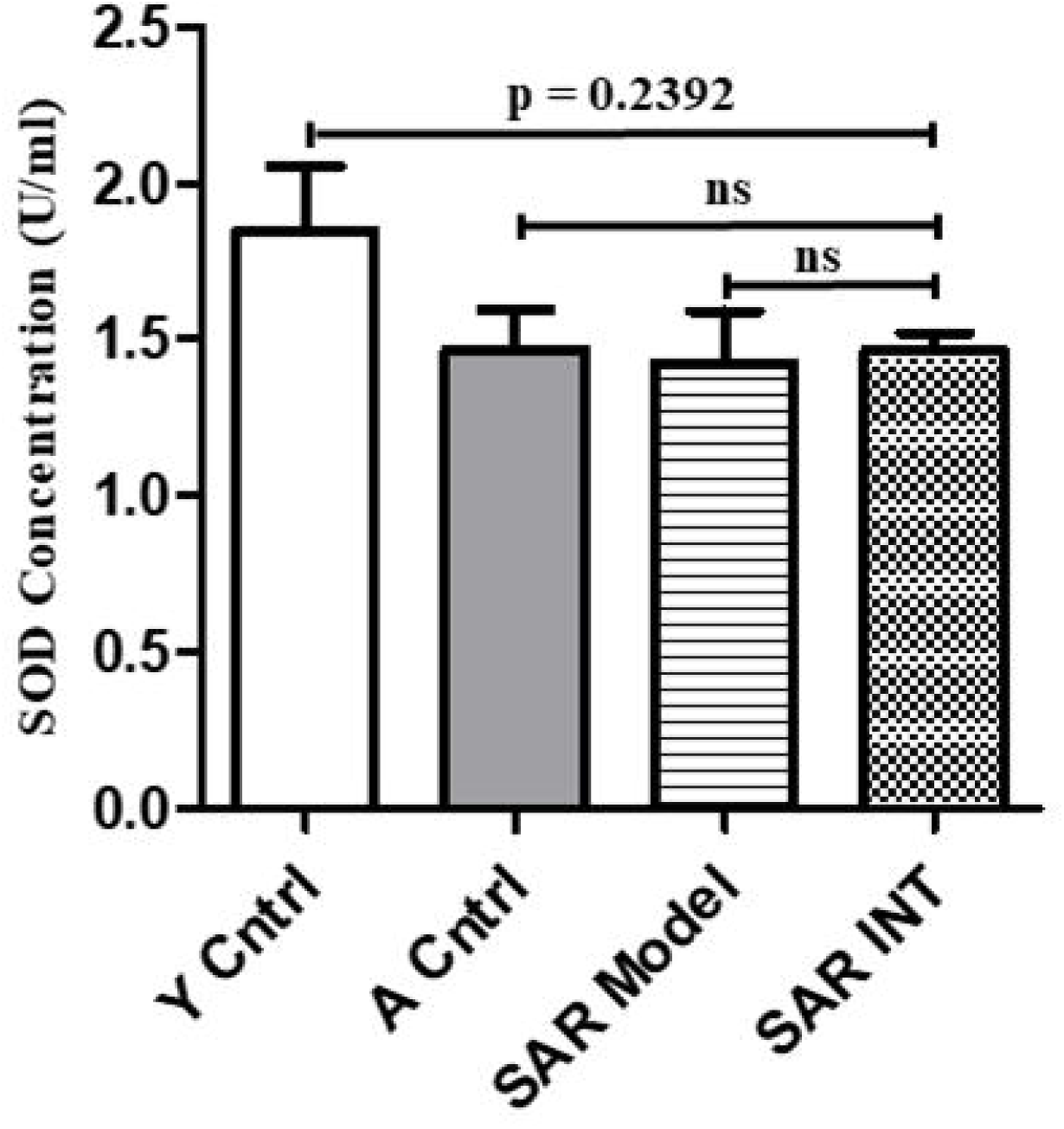
Differential expression of the SOD enzyme in Y Cntrl, A Cntrl, SAR Model and SAR-INT (n =3).

## 3. Discussion

Animal models to explore the progression of the loss of skeletal muscle mass and strength do not exist (Palus et al., 2017). Thus, it is imperative to develop an experimental animal model to determine the perturbations at the genomic or proteomic level that occur at the onset of sarcopenia. In this study, we assessed animals under both sedentary confinement and dietary protein restriction to resemble a sarcopenic ageing phenotype. Physical parameters, namely, body weight, grip strength and gait speed, were assessed to demonstrate functional deficits of sarcopenia. Mitochondrial oxidative enzymes were assayed in muscle tissue to evaluate the possible role of oxidative stress in the development of the sarcopenic state.

Changes in animal body weight have been found to proportionally increase with age (Gargiulo et al., 2012). Our data revealed that the body weight of the A-Cntrl group was considerably higher compared to that of the Y-Cntrl group, most probably due to the accumulation of fat mass, which is consistent with a previous report (Yang, Smith, Keating, Allison, & Nagy, 2014)). However, an inverse correlation was found for the SAR-model group, which phenotypically explains the importance of dietary intake of essential amino acids. The decline in the body mass of the SAR-model group suggested the presence of additional parameters of sarcopenia in the experimental animals. Diet regulates the body weight of animals during the process of ageing (Yang et al., 2014). The animals in this study were subjected to grip strength analysis an established parameter for sarcopenia in humans. Sternäng and colleagues provided a detailed description of the decline in human grip strength (Sternäng et al., 2015). Age-associated decreases in grip strength and muscle mass in mice have been assessed and are well correlated with tumour suppressor genes in MuSCs (Sousa-Victor et al., 2014). Our results for the grip strength analysis showed a correlation with age (A-Cntrl); however, compared with the A-Cntrl group, the SAR-model group showed a 30% decline in grip strength. Grip strength value of N/kg< 9 was considered as the threshold for sarcopenia in the SAR-model group. The recently described criteria for the diagnosis of sarcopenia include low muscle mass, which is an important causative factor, followed by low muscle strength and poor physical performance (Cruz-Jentoft et al., 2010; Han, Bokshan, Marcaccio, DePasse, & Daniels, 2018). Animals with grip strength force <9 N/kg underwent further physical performance assessment. Voluntary wheel running by a mouse is considered a reliable technique to assess physical performance and endurance (Goh & Ladiges, 2015). Voluntary wheel running in this study showed decreased physical performance in the SAR-model group compared with the A-Cntrl group. The SAR-model group exhibited a longer elapsed time (in seconds) than the A-Cntrl group, showing a 28% increase in the time required to cover a distance of 100 m. This result shows the low physical performance of the SAR-model group, which may be due to the dietary and mobility restrictions imposed on the animals. Sedentary lifestyle due to low physical activity leads to functional decline (Bogdanis, 2012).The low protein diet leads to a negative balance (Moore, 2014) between muscle protein synthesis and muscle protein breakdown (Moore et al., 2015). Thus, all these aspects contributed to the development of sarcopenia.

The tissue-specific decline in skeletal muscle mass was quantified by recording the wet weight of three different muscles: TA, EDL and GAST. The weights of TA and GAST were significantly decreased in the SAR-model group compared to the A-Cntrl group. These observations suggested that the cause of the decreased body weight was the restrictive conditions. The oxidative capacities of three different mitochondrial enzymes (ATPase, NADH-TR and SDH with COX staining) were assessed in TA tissue. The ATPase staining of type II fibres at the basic pH of 10.8 revealed a lower intensity in the SAR-model group than in the A-Cntrl group due to low ATPase enzymatic activity. Additionally, the CSA of type II fibres decreased in our study, in accordance with previously reported data on aged mice (Frontera et al., 2000).The observed weak staining of NADH-TR and SDH enzymes with COX revealed that mitochondrial enzymatic activity was decreased in the SAR-model group compared with the A-Cntrl group. The causative factor for this result could be catabolic stress and muscle wasting due to the lack of essential amino acids. Taken together, these results clearly emphasize the importance of mitochondrial enzymatic activity for skeletal muscle health and suggest that these enzymes can be considered important therapeutic targets to attenuate the age-related deterioration of muscle function and metabolism.

The role of MuSCs in the SAR-model group and the impact of interventions on the MuSC niche were assessed by isolating and quantifying these cells. The small MuSC populations in the A-Cntrl group are consistent with the previous findings of Victor and colleague (Sousa-Victor et al., 2014). Compared with the A-Cntrl group, the SAR-model group had significantly fewer Ki67-positive cells MuSCs, although there was no significant difference in the overall MuSC population. This observation indicates a significant decline in the MuSC proliferation capacity of SAR-model animals, possibly due to the increased expression of tumour suppressor genes or excessive load of oxidative insult to MuSCs in these animals compared with the control animals.

ROS-induced oxidative stress plays a crucial role in sarcopenia, as oxidative stress is one of the causative factors for the apoptosis of progenitor and mature skeletal muscle cells (Bellanti et al., 2018; Weindruch & Sohal, 1997).It has also been reported that physical exercise induces the up-regulation of antioxidant defence in skeletal muscle (Ji, 2007, Gaetano Santulli et al., 2013) and antioxidant gene expression (Gomez-Cabrera et al., 2008). It has been reported in the literature that skeletal muscle activity improved by increasing the mitochrondrial antioxidant enzyme and also by voluntary exercise in aged mice (Xie et al., 2015). ROS production increases in ageing which dysregulates mitochrondrial homeostasis due to deficiency of cellular antioxidant defense in sarcopenia (Lopez-Otin et al., 2013). The resulting oxidative stress affects the mitochrondrial protein function (Baraibar et al., 2013) and weakens the muscles. Exercise, on the other hand, benefits mitochondrial function by activating mitochondrial biogenesis and mitophagy (Romanello & Sandri 2016).Exercise improved the metabolic potential of muscle by increasing oxidative capacity (Santulli et al., 2013).

Sestrin1 and sestrin2 are over-expressed in the skeletal muscles (Velasco-Miguel, et al., (1999). Sesn are regulated by p53, and serum p53 increases after vigorous exercise (Budanov, 2011). Lack of SESN increases ROS and thereby promotes oxidative stress (Budanov, A.V, et al., 2004). Loss of serum sestrin in frail elderly compared to non-frail elderly was reported in our earlier study (Rai, et 2018). Physical exercise promotes the intracellular ATP depletion and the increase of AMP content, triggering AMPK activation. Another classic effect of physical exercise is the increase of antioxidant activity by inducing anti-oxidant enzymes (Barbosa et al., 2012). There is evidence that Sesns increase the effect of physical exercise during oxidative stress and activate AMP, thus strengthening the muscle /mass by improving insulin sensitivity and autophagy induction in old mice (Lenhare, et al., 2017; Barbara et al., 2018). Current research is directed at understanding the cellular and sub-cellular basis of human sarcopenia derived from the earlier knowledge. Other reports have also explained increased level of ROS generation and enhanced antioxidant enzymes due to regular exercise (Lucia et al 2013).

. Considering oxidative stress conditions, we explored antioxidant Sestrins in our animal model. Sestrins have been studied for their role in skeletal muscle. Sestrins, as antioxidant molecules, show altered expression in frail older patients and play a protective role under stressful conditions (Rai et al., 2018). The present study suggests that reduced expression of Sestrins can be one of the causative factors responsible for sarcopenia in the SAR-model group, which failed to cope with oxidative stress. Compared with the SAR-model group, the SAR-INT group exhibited a 27.6% increase in Sestrin2 concentration as well as SOD level. It can be assumed that the high expression of Sestrin2 in the SAR-INT group was due to physical activity which regulated the antioxidant defence system in sarcopenia.

Sarcopenia is an important clinical challenge for ageing population globally. With uncertainty in understanding its origin, no interventionis currently available to reverse sarcopenia. The animal model developed in the present study by protein-deficient diet with a forced sedentary lifestyle we created a novel model similar to human sarcopenia. Physical activity appears to be a simple intervention that can improve sarcopenia, as was observed in the SAR-model animals modulated by antioxidant molecules. This study also indicates a role for oxidative stress in the development of sarcopenia. The role of the antioxidant proteins Sestrins may play a promising role for the clinical assessment of sacropenia in the future.

## 4. Experimental Procedures

### Animals

Central Animal Facility of All India Institute of Medical Sciences, New Delhi provided eighty 6-to 8-week-oldmale C57BL/6 mice for the study. All animals were fed *ad libitum* (Ibebunjo et al., 2013)with rodent maintenance (RM) diet for the first 14 months. The animals were housed in plastic cages at a temperature of 23±2°C and a relative humidity of 55±15% and under a 12-hour light and 12-hour dark cycle. A group of animals (n = 5) was sacrificedat this point to serve as the Y-Cntrl group. The remaining animals were divided into two groups:

The first group, A-Cntrl, was allowed to age with an *ad libitum* diet and was sacrificed at 21 months. The second group was maintained on a protein-deficient diet with a forced sedentary lifestyle in a small cage to develop sarcopenic model (SAR-model). Thirty animals were assigned to the SAR-model group and were individually housed in different (specifically modified) cages of half the length of a standard small cage (26×10×13 cm). A confinement with the dimensions13×10×13 cm was designed inside the standard cage to provide sedentary isolation (Figueiredo et al., 2009).The model animals were kept in the confinement cage for the defined time period of seven months. The animals were fed a customized 14% protein-deficient diet procured from Teklad Global (2014S, Teklad Global, USA). At 21 months, this group was subjected to physical activity intervention (SAR-INT). The SAR-INT animals were sacrificed at 24 months. Thus, for various post-mortem experiments, four animal groups were created:

i. Young (Y-Cntrl) - 14 months
ii. Aged control (A-Cntrl) - 21 months
iii. Sarcopenic Protein-Deficient model (SAR-model) - 21 months
iv. Post-intervention SAR-model(SAR-INT) - 24 months

Animal experiments were approved by the Institute Animal Ethics Committee of All India Institute of Medical Sciences, New Delhi.

### 4.1 Physiological parameter Tests

For the phenotypic characterization of all animals from different groups, physical assessment was performed before sacrifice. The parameters for assessment were body weight, grip strength and the time required to run in an activity wheel.

#### 4.1.1 Body weight measurement

The body weight of each animal was recorded twice before the animal was sacrificed (Palmer et al., 2009). The mean body weight of each group was calculated to analyse the differences among ages and interventions.

#### 4.1.2 Grip Strength

A grip strength meter (IITC Life Science, USA) was used to measure the muscle strength of the animals. The apparatus consisted of a digital display connected to specifically designed forelimb grasping pull bar made of steel. Uniform gentle force was applied by holding the tail of the animal, while the animal tried to pull back by gripping the T-rod bar until the grip was released. The maximum strength value to release the T-rod bar was displayed on the screen and recorded as grip strength. Using the above methodology, grip strength was measured after allowing the animal to become familiar with the grip strength meter for half an hour as the acclimatization time. Six trials were successively performed with a rest interval of 2 minutes for each animal in all fourgroups (Nagayach, Patro, & Patro, 2014).

#### 4.1.3 Time elapse analysis

Voluntary wheel running was used for two purposes: to test the endurance capacity and as exercise for the intervention group. The experimental activity wheel was 15 cm in diameter. The number of cycles and the time required by the animals were recorded by the digital display system connected to the activity wheel. The animals in the Y-Ctrl, A-Ctrl, and SAR-model study groups were subjected to a wheel running test to analyse endurance capacity. The time in seconds required to cover a distance of 100 metres was recorded. To establish the SAR-INT group, the SAR-model group animals were allowed to voluntarily run on the wheel for physical activity for a period of 10 weeks after the initial assessment SAR-model group, and the difference in time elapsed to cover the distance of 100 metres was assessed. A fixed number of revolutions (n = 210) was set for each animal.

#### 4.1.4 Intervention regime by *Activity Wheel test*

An intervention protocol for voluntary wheel running was developed for the SAR-INT group to assess the health-enhancing benefits of exercise. The activity wheel running protocol was administered five days per week for a period often weeks (Goh & Ladiges, 2015). A fixed number of revolutions (n = 210) was set for each animal. The time required to complete a 100-m run was digitally recorded.

### 4.2 Histology, morphometry and enzyme histochemistry

Animals were sacrificed using a mixture of ketamine and xylazine (80 mg: 10 mg/kg).The skeletal muscles from both limbs were dissected and used for various measurements and experiments.

#### 4.2.1 Wet tissue weight analysis

The skeletal muscle tissues were isolated from sacrificed animals. TA, EDL, and GAST were rinsed in PBS, and the wet weights were recorded. Samples were stored at −80°C for subsequent experiments (Hahn et al., 2009). The mean tissue weight was calculated to assess tissue-specific changes in all four groups.

#### 4.2.2 Histological staining with mitochrodrial enzymes

The TA muscle was snap-frozen in 2-methylbutaneprein liquid nitrogen. Serial8-µm-thick cryosections were cut, mounted on lysine-coated glass slides (Sigma, USA),air dried and stored at −80°C (Gouspillou et al., 2014). Cryosections were processed and examined for the in situ determination of fibre type composition, CSA and mitochondrial oxidative capacity of the skeletal muscle tissue. Sections were stained with haematoxylin and eosin to assess the general tissue architecture of myofibres using a standard protocol.

Sections were stained for ATPase (pH 10.5) activity according to an established protocol (Guth & Samaha, 1969; Suga et al., 2011). Sections stained for ATPase activity were used to classify the fibres as type I or II and to estimate the relative ATPase expression. ATPase at pH 10.8 distinguishes type I (light staining) from type II (dark staining) fibre. Sections were incubated for 15 minutes at pH 10.8 followed by incubation in 20 mM sodium barbital at pH 9.5 with 9 mM CaCl_2_ and 2.7 mM ATP for 45 minutes. Subsequently, the samples were rinsed twice in 1% CaCl_2_ (1 minute each wash), immersed for 2 minutes in 2% CaCl_2_ and rinsed several times with tap water. After staining with 1% (NH_4_)_2_S, the sections were washed several times with tap water, dehydrated with ethanol, cleared in xylene, and mounted.

Routine NADH-TR staining was performed as previously described (Brooke & Kaiser, 1970; Hämäläinen & Pette, 1993) to semi-quantitatively evaluate the oxidative capacity of the fibres. Briefly, a 1:1 mixture ofnitro-blue tetrazolium (NBT) solution and NADH solution was added to muscle sections and incubated for 30 minutes at 37 °C. The incubated sections were washed three times with tap water, washed with an increasing gradient of three solutions of 30%, 60% and 90% acetone-deionized water, and finally washed with a decreasing gradient of acetone-deionized water solutions to remove the unbound NBT. The sections were rinsed three additional times with tap water, mounted with DPX mountant, air dried, and stored at room temperature. Cryostat sections were also subjected to the sequential histochemical staining of COX/SDH as previously described (Díaz-Herrera et al., 2001). Briefly, sections were incubated for 45 minutes at 37 °C with COX reaction solution and for 40 minutes at 37 °C with SDH solution (pH 7.0).

### 4.3 Double Immunofluorescence Assay

Double immuno-staining was performed by sequentially adding primary and secondary antibodies. Cryostat sections of TA tissues were pre-fixed with ice cold methanol stored at −80 °C. Sections were acclimatized to room temperature, followed by washing with PBS. After blocking at room temperature for 1 hour, the sections were incubated for 90 minutes with the primary antibodies Pax 7 (monoclonal DSHB, USA) and laminin, (Bio Legend, USA) at dilutions of 1:200 and 1:500, respectively. After washing with PBST (3 times for 5 minutes each), the tissue sections were incubated with secondary antibodies (1:200, Alexa Fluor 594/AlexaFluor647,BioLegend, USA) for 60 minutes. The slides were mounted with Vecta-shield mounting medium containing DAPI (Invitrogen, USA) to label nuclei. Fluorescence images were acquired using a Zeiss Imager M1 Fluorescence microscope.

### Dissociation of Skeletal muscle and single cell preparation

Sacrificed animals were used to isolate whole skeletal muscle tissue from both limbs. Dissected skeletal muscle tissue was immediately subjected to mincing following the manufacturer’s protocol to form a smooth pulp using a skeletal muscle tissue dissociation kit (Cat no.130-098-305, Miltenyi Biotec, Germany). Dissociated tissue was filtered using a 70-µm filter (Mitenyi Biotec, Germany) to remove the cell clump from the single cell suspension.

### 4.4 MuSCs Isolation and Quantification

Whole skeletal muscle tissue from both limbs of the animals from all groups was dissected. Mincing was immediately performed following the manufacturer’s protocol to form a smooth pulp using a skeletal muscle tissue dissociation kit (Miltenyi Biotec, Germany). The dissociated tissue was filtered using a 70-µm filter (Mitenyi Biotech, Germany) to remove the cell clumps from the single cell suspension. MuSCs were isolated from the single cell suspension using the MACS technique. A satellite cell isolation kit was used to separate MuSCs according to the manufacturer’s instructions (Cat no. 130-104-268, MiltenyiBiotec, Germany). The isolated cells were further enriched using anti-alpha integrin 7 micro-beads (Miltenyi Biotec, Germany) to obtain a pure population of cells.

#### 4.4.1 Flow Cytometry Analysis

The enriched population of cells was processed for double staining flow cytometric analysis (Attune Nxt Flow Cytometer, Invitrogen, USA). The cells were incubated with primary antibodies at a titrated dilution for 30 minutes at 4°C in supplemented PBS containing 2 mM EDTA and 2% FBS at ∼1 million cells per ml. Phycoerythrin-conjugated anti-alpha 7-integrin antibody (130-102-716 Miltenyi Biotec, Germany) was used as a marker for MuSCs, and FITC-conjugated anti-Ki 67 (BioLegend, USA) was used as a proliferation marker.

### 4.5 Surface Plasmon Resonance (SPR)

The differential expression profile of Sestrin family proteins in cell lysate was determined by SPR technology using the BIACORE 3000 system (Wipro GE Healthcare, UK). The quantification of Sestrin1 and Sestrin2 levels was performed in all four experimental groups. This technique provides a unique label-free platform for the analysis of specific bimolecular interactions in realtime. A CM5 sensor chip (Wipro GE Healthcare, UK) was used to immobilize the antibodies via goat anti-mouse Sestrin1 IgG (SC-376170, Santa Cruz, USA) and rabbit anti-mouse Sestrin 2 IgG (SC-292558, Santa Cruz, USA). Immobilization was performed using the amine coupling reaction kit (Wipro GE Healthcare, UK) in separate flow cells of the chip. The protein samples were dissolved in 20 mM sodium acetate (pH 4.5) at a concentration of 10 μg/ml and were allowed to run at a flow rate of 5 μl/min. BIA evaluation software version 4.1 was used to evaluate the results.

### 4.6 Western blot analysis

MACS-isolated MuSCs were suspended in ice cold lysis buffer and homogenized using a gentle MACS dissociator (Miltenyi Biotech, Germany) to extract the cell lysate. The homogenate was centrifuged at 12000 rpm for 15 min at 4°C to isolate total supernatant protein. Protein concentration was determined with a Pierce BCA Protein Assay Kit (Thermo Scientific, USA) according to the manufacturer’s instructions. The lysate was filtered and stored at −80 °C till use. The protein extracted from mouse MuSC lysates was separated using SDS-PAGE and transferred to polyvinylidenedifluoride (PVDF) membranes. The membrane was blocked using 5% non-fat milk. The PVDF membrane was washed with TBST. Antibodies against Sestrin1 and Sestrin2 were incubated overnight at dilutions of 1:200 and 1:400, respectively. After washing, a one-hour incubation with secondary horseradish peroxidise (HRP)-conjugated goat anti-mouse IgG and rabbit anti-mouse IgG (1:4000) was performed. Protein expression was visualized using an enhanced chemiluminescent detection system (Thermo Scientific, USA). The relative and normalized protein expression was calculated based on GAPDH expression. Band density was evaluated by My Image Analysis Software (Life Technologies, USA).

### 4.7 SOD activity

SOD activity was measured by a commercially available enzyme activity kit according to the manufacturer’s instructions (Cayman Chemical, Ann Arbor, Superoxide Dismutase Assay Kit; 706002). Briefly, samples were homogenized 1:10 (mgwt/μL buffer) using gentle macs dissociator (Cat no. 130-093-235, MiltenyiBiotec, Germany) in 20mM HEPES, 1mM EGTA, 210mM mannitol, and 70mM sucrose (pH7.2) and centrifuged at 1500g for 5min at 4°C. SOD activity obtained was normalized to protein concentration in the lysate.

### Data Analysis

Statistical analysis was performed using Graph Pad Prism 5.0 software, and p<0.05 was considered statistically significant. To compare variables, paired and unpaired t-tests between young and aged group and one-way analysis of variance (ANOVA) were performed between all four study groups mentioned above with Tukey’s Multiple Comparison post hoc tests.

## Acknowledgement

AIIMS, New Delhi, provided fellowship to the first author and funds for procurement of equipment and consumables.

## Author Contributions

ABD and SKP provided the study concept and design; data analysis and preparation of the manuscript. MA performed the experiments and prepared the manuscript. SD plannedtheexperiments and assisted in manuscript preparation. SRM guided the flow cytometry experiments and helped in data analysis. DP provided study resources, and gave valuable suggestions.SS helped in SPR methods. None of the author had any conflict of interest.

## Supplementary Figures

**Figure:** S-1 Difference in cross sectional area of type II fibers in different experimental groups Y Cntrl, A Cntrl, SAR Model and SAR-INT.

**Figure:** S-2 Hematoxylin and eosin staining of skeletal muscle tissue showing intact morphology before immunofluorescence.

