## Supplementary figures and images for "Physical activity improves sarcopenia in a murine model by enhancing the proliferative potential of muscle stem cells, oxidative capacity of mitochondrial enzymes and expression of Sestrins"

### Supplementary material Fig S1

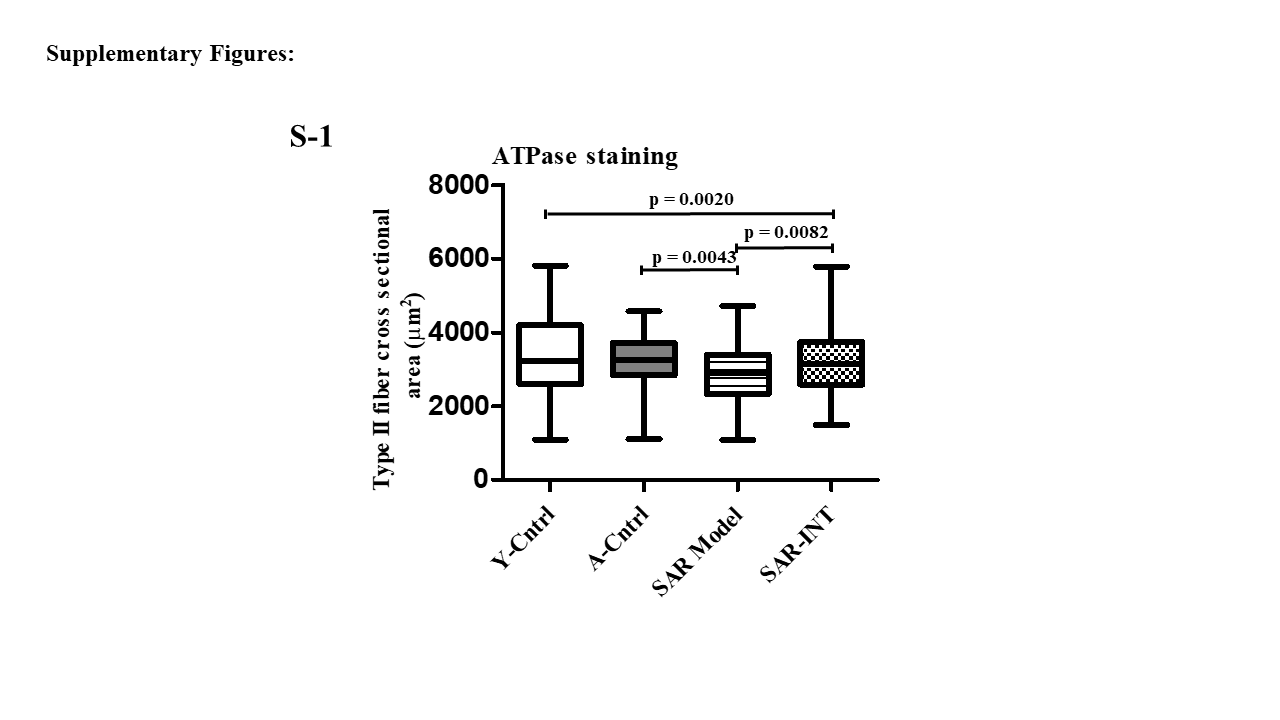

### Supplementary material Fig S2

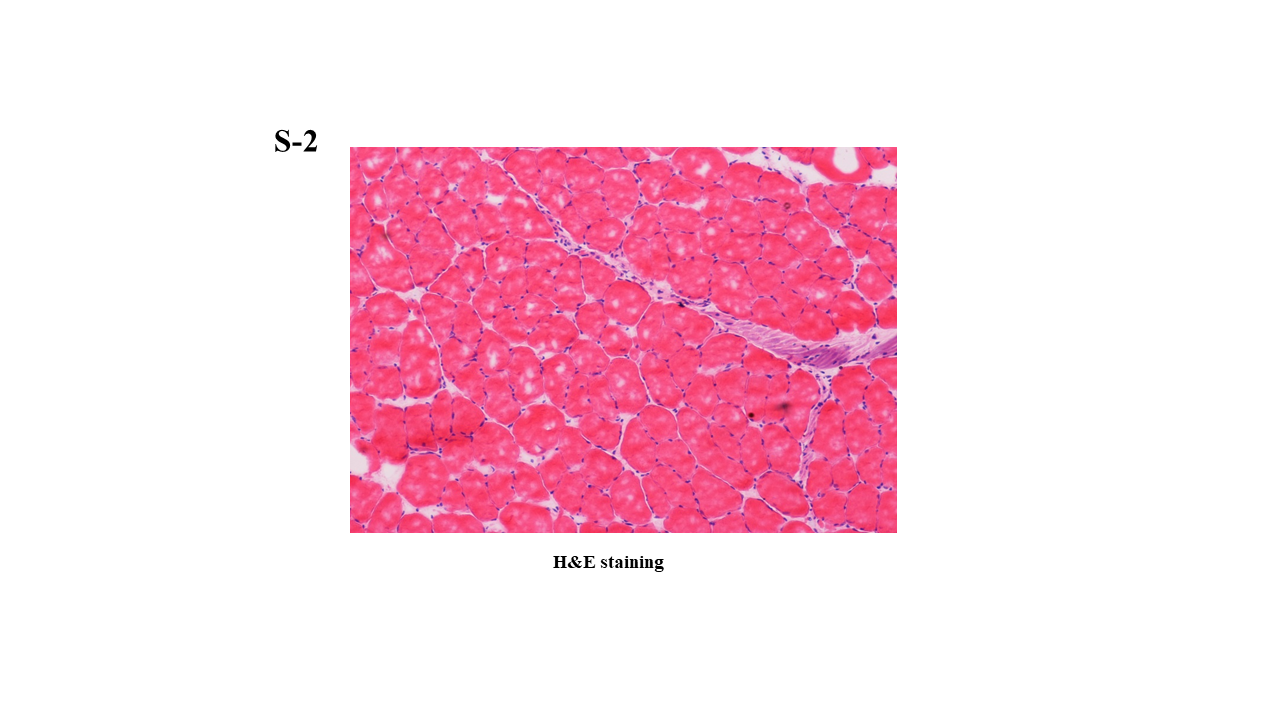
